# Prediction of molecular markers of bovine mastitis by meta-analysis of differentially expressed genes using combined p-value and robust rank aggregation

**DOI:** 10.1101/2021.11.24.469841

**Authors:** Anushri Umesh, Praveen Kumar Guttula, Mukesh Kumar Gupta

**Affiliations:** Department of Biotechnology and Medical Engineering, National Institute of Technology Rourkela, Odisha 769008, India; Bioinformatics Scientist, Department of Clinical Hematology, Tata Medical Center, Kolkata 700160, India; Center for Bioinformatics and Computational Biology, National Institute of Technology Rourkela, Odisha 769008, India

**Keywords:** Mastitis, meta-analysis, combined p-value method, Rank aggregation method, miRNA-Gene targets, mammary epithelial cells

## Abstract

Bovine mastitis causes significant economic loss to the dairy industry by affecting milk quality and quantity. *E.coli* and *S.aureus* are the two common mastitis-causing bacteria among the consortia of mastitis pathogens, wherein *E.coli* is an opportunistic environmental pathogen, and *S.aureus* is a contagious pathogen. This study was designed to predict molecular markers of bovine mastitis by meta-analysis of differentially expressed genes (DEG) in *E.coli* or *S.aureus* infected mammary epithelial cells (MECs) using p-value combination and robust rank aggregation (RRA) methods. High throughput transcriptome of bovine (MECs, infected with *E.coli* or *S.aureus*, were analyzed, and correlation of z-scores were computed for the expression datasets to identify the lineage profile and functional ontology of DEGs. Key pathways enriched in infected MECs were deciphered by Gene Set Enrichment Analysis (GSEA), following which combined p-value and RRA were used to perform DEG meta-analysis to limit type I error in the analysis. The miRNA-Gene networks were then built to uncover potential molecular markers of mastitis. Lineage profiling of MECs showed that the gene expression levels were associated with mammary tissue lineage. The up-regulated genes were enriched in immune-related pathways whereas down-regulated genes influenced the cellular processes. GSEA analysis of DEGs deciphered the involvement of Toll-like receptor (TLR), and NF- Kappa B signalling pathway during infection. Comparison after meta-analysis yielded with genes ZC3H12A, RND1 and MAP3K8 having significant expression levels in both *E.coli* and *S.aureus* dataset and on evaluating miRNA-Gene network 7 pairs were common to both sets identifying them as potential molecular markers.

## 1. Introduction

Mastitis, the inflammation of the mammary gland, is the most common disease in cows with various adverse consequences on animal health and farm economics. Bovine mastitis has been estimated to cause a loss in gross profit of USD 516-722 per cow to the farmers (Puerto et al. 2021). Bacterial toxins and tissue damage due to host reactions in mastitis also alter the milk quality that can severely affect the health of animals, neonates, and consumers (Kaliwal 2013). Consequently, milk producers and veterinary professionals have been consistently looking for effective and economical ways to prevent and treat mastitis in bovine herds. Diagnosis of mastitis is usually made by tests such as Somatic Cell Count (SCC) (Alhussien and Dang 2018) and the California Mastitis Test (CMT), which are based on the presence of somatic cells and leukocytes in the milk (Ferronatto et al. 2018). However, these methods are non-specific and may produce false positives due to conditions such as stress experienced by animals (Liu et al. 2019). Thus, there is a need to develop newer molecular markers that can non-invasively predict and identify bovine mastitis with minimal errors and high specificity.

Bovine mastitis is usually caused by bacterial pathogens, which can be broadly grouped into environmental pathogens (e.g., *Escherichia coli*, *Enterobacter faecalis*, *Klebsiella* spp., *Enterobacter* spp., *Citrobacter* spp., *Streptococcus* spp. etc.) and contagious pathogens (e.g., *Staphylococcus aureus*, *Streptococcus agalactiae*, *Corynebacterium bovis*, *Mycoplasma bovis*, etc.) (Reshi et al. 2015; Dufour et al. 2019). Among the consortia of mastitis-causing bacteria, *E.coli* and *S. aureus* are the two most common pathogens that are Gram-negative and Gram-positive, respectively (Schukken et al. 2011). The Gram-negative bacteria are known to trigger a more severe and acute host response, with more intense innate immune reactions, than Gram-positive bacteria (Kusebauch et al. 2018). Although the pathogenesis and host reaction to mastitis is not entirely understood, there has been evidence of differences in gene and protein expression levels that specifically differ for different infecting pathogens (Kusebauch et al. 2018). Several proteins’ expression levels were reported to be upregulated in mastitis, and include cytokine-cytokine receptor interaction pathways that produce chemotactic signals to attract infiltration of neutrophils to the inflammatory site for killing the pathogens (Ziyin Han, Yongliang Fan, Zhangping Yang, Juan J. Loor 2020). A significant change has also been reported in the milk metabolites, with infected groups showing elevated phenyl pyruvic acid and homogentisic acid (Tong et al. 2019). Thus, genes and proteins involved in various signalling pathways affected in mastitis may serve as possible molecular markers for detection of bovine mastitis.

MicroRNAs (miRNAs) play essential roles in the post-transcriptional regulation of gene expression by binding and degrading the target mRNA transcripts (Sun et al. 2019). The milk and mammary gland also contain a plethora of miRNAs, whose expression profile changes during the inflammatory response to pathogens (Cintio et al. 2020). Differences in the expression level of miRNAs and other non-coding RNAs (ncRNAs) have been reported between mastitis-affected and unaffected animals (Lai et al. 2020). The miRNA-Gene target networks involving differentially expressed genes (DEGs) targeted by miRNAs can also be the potential molecular markers for the identification of mastitis (Islam et al. 2020). However, individual studies may provide biases based on animal demographics and the physiological condition of the animals at the time of the study (Burvenich et al. 2007). A meta-analysis of the dataset increases the statistical power and reduces the individual variability present in datasets (Hong and Breitling 2008). Effect size combination, rank combination and p-value combination being the three methods for meta-analysis with p-value combination providing the advantage over others by its extensibility to combination of studies with variable outcome and genomic study standardization across a common scale in detection of novel markers (Tseng et al. 2012).

Thus, in this study, a meta-analysis of high throughput transcriptomic data from mammary epithelial cells (MECs), infected with *S.aureus* or *E.coli*, was carried out to identify potential molecular markers of bovine mastitis. The transcriptomes of MECs were chosen for the study because MECs can be non-invasively collected from the milk (Buehring 1990). Moreover, unlike whole milk or mammary tissue, analysis of MECs reduces the variability factors from blood or other cell types present in the tissue (Buehring 1990). Two meta-analysis methods were compared to identify significant DEGs of mastitis. Rational behind using two methods in the study was to decrease the false positives and to understand the difference in results obtained by comparing two meta-analysis methods. Gene Set Enrichment Analysis (GSEA) was performed to decipher the molecular pathways that were affected in the infected MECs. The miRNA-Gene targets were then predicted to identify potential molecular markers of bovine mastitis.

## 2. Methodology

Identification of molecular markers of mastitis through meta-analysis of DEGs followed a sequential process of data collection, pre-processing and functional analysis of transcriptomes. A brief overview of the methods followed in the study is illustrated in Fig 1.

**Fig. 1:**
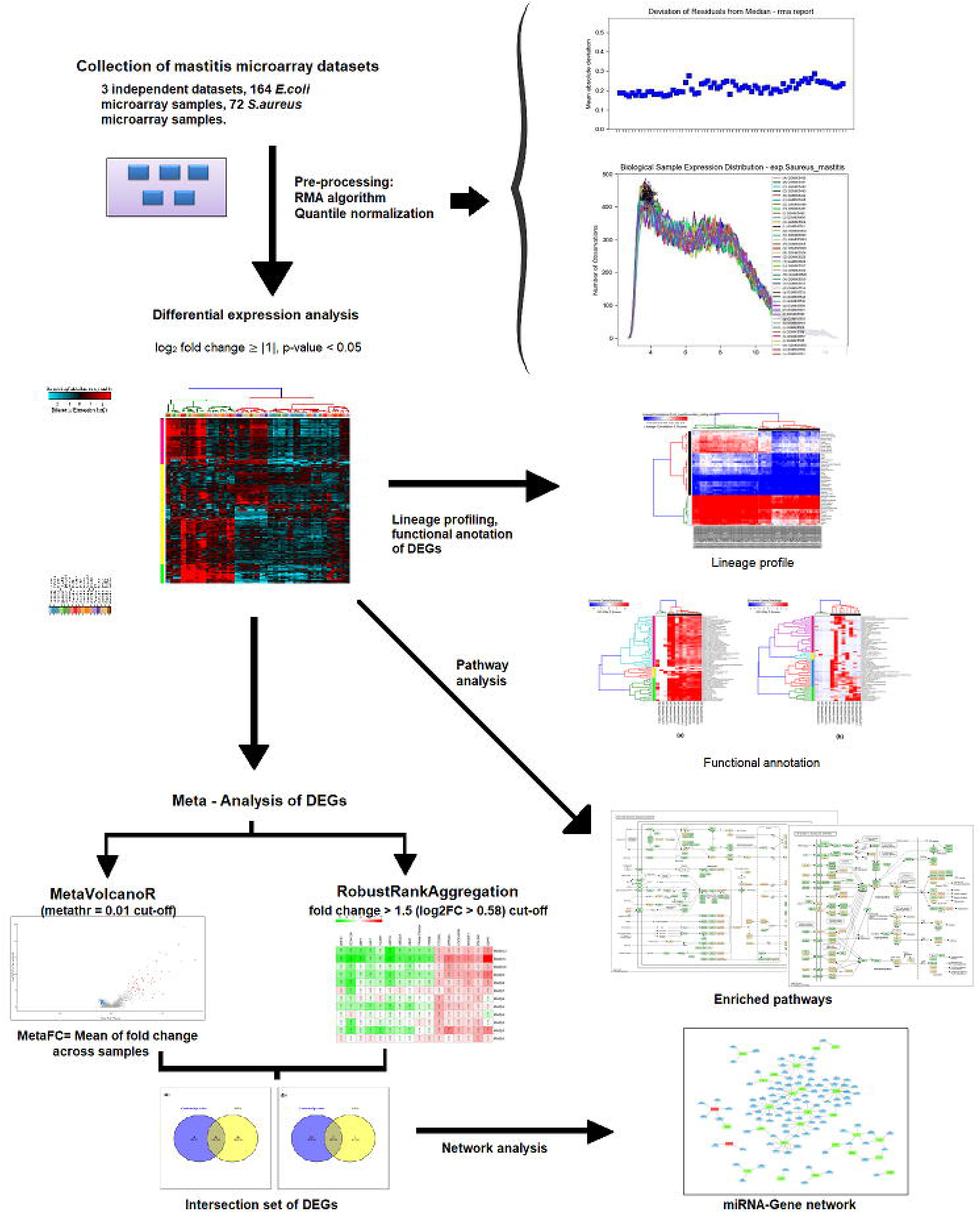
Overview of the methodology used in this study.

### 2.1 Datasets

Datasets for DEG analysis were retrieved from Gene Expression Omnibus of the National Center for Biotechnology Information (URL: https://www.ncbi.nlm.nih.gov/geo/). The keywords “mastitis”, “transcriptome”, “mRNA”, “*Bos taurus*” and “bovine” were used in advanced search options. Microarray datasets belonging to transcriptome analysis of bovine MECs on the Bovine Affymetrix array platform were manually selected for the study. The *E.coli* study included three datasets *viz*. GSE24560, GSE25413, GSE32186, whereas the *S.aureus* study included two datasets *viz*. GSE24560 and GSE25413. The GSE24560 consisted of 27 controls with 31 and 30 samples of *E.coli* and *S.aureus* infected MECs, respectively. The GSE25413 had six controls and 12 each of *E.coli* and *S.aureus* samples, respectively. The GSE32186 dataset consisted of 6 controls and 6 *E.coli* infected MEC samples. In total, 85 datasets of *E.coli* and 72 datasets of *S.aureus*-infected MECs were included in this study.

### 2.2 Pre-processing of data and identification of DEGs among control and infected MECs

Pre-processing of GSE datasets and identification of DEGs among healthy control and infected MECs were performed using AltAnalyze software (Emig et al. 2010), as we described earlier (Huma et al. 2020) Briefly, the. CEL files were subjected to pre-processing and Robust Multi-Array Average (RMA) algorithm for background correction and normalization of gene expression data from microarray analysis (Irizarry et al. 2003). Quantile normalization was applied to the processed data by which the probe intensity distribution was made the same across all GeneChip arrays before further quantification. The group comparison was carried out using moderated t-test, and the Benjamini-Hochberg method was used for adjusted p-value calculation (Islam et al. 2020). GO-Elite was used for analyzing the gene ontologies with gene having log 2 fold change ≥ |1| and p-value < 0.05 as a cut-off value, as described earlier (Han 2019; Li et al. 2019). Pruning of ontology terms was done using z-scores, and over-representation assay for the DEGs was carried out using Fisher’s exact test (Emig et al. 2010). Z-score was calculated by dividing the normal distribution by the hyper-geometric distribution.

### 2.3 GSEA of DEGs among control and infected MECs

The GSEA of DEGs among MECs from healthy controls and those infected with *S.aureus or E.coli* was analyzed using WebGeStalt (Liao et al. 2019). Genes differentially expressed in all the analyzed datasets were subjected to the GSEA. The inputs were given as gene lists with their corresponding fold changes obtained from AltAnalyze after performing DEG analysis. Top positively and negatively enriched pathways were studied if they were significant with a false discover rate (FDR) of < 0.05. Enrichment of genes significant in *E.coli* and *S.aureus* infected MECs was calculated by comparing significant genes with reference gene set proportion to the gene included in a given category.

### 2.4 Meta-analysis of DEGs among control and infected MECs

Meta-analysis of the DEGs from all studies was done using the combined p-value method and robust rank aggregation (RRA) method, as described below. These two methods were selected based on the criteria that the conditions of carrying out the experimental analysis varied, and the rank combination and p-value combination are better suited for meta-analysis of these data in comparison to effect size combination (Chang et al. 2013). An intersection set of DEGs obtained from the two meta-analysis methods was created using Venny 2.1 (https://bioinfogp.cnb.csic.es/tools/venny/index.html), and the DEGs reported by both methods as significant were used for the construction of the miRNA-Gene network.

#### 2.4.1 Combined p-value method for meta-analysis

The combined p-value for meta-analysis of gene expression data was performed in MetaVolcanoR, as described earlier (Prada C, Lima D 2020). Briefly, the Excel files containing all the datasets for each of the samples were prepared. The input file consisted of the Gene symbols with their corresponding log2 of fold changes and adjusted p-values from the analysis. MetaVolcanoR was used for obtaining the significant genes from the combined datasets, and the screening was done based on a metathr cut-off of 0.01. The metaFC was calculated by taking the mean of the repeated values in the gene expression dataset.

#### 2.4.2 RRA method for meta-analysis

The RRA method for meta-analysis of DEGs was performed in R, as described earlier (Kolde et al. 2012). Briefly, the differential gene expression data, obtained from AltAnalyze, were first divided into two rank matrices: up-matrix and down-matrix, respectively. Rank aggregation was then applied to these matrices, and the p-values were adjusted using the Bonferroni method for statistical significance. Genes having fold change of ≥ |1.5| were considered significant and used to construct the miRNA-Gene network (Swanson et al. 2009; Islam et al. 2020).

### 2.5 miRNA-Gene network of DEGs among control and infected MECs

Up- and down-regulated genes obtained from the intersection of two meta-analysis methods were submitted to the miRNet analysis platform, and the potential miRNA-Gene targets were obtained (Chang et al. 2020). The miRNA-Gene targets were predicted by the miRanda algorithm (Enright et al. 2003), and only significant interactions were screened and given as output in the platform. Algorithm miRanda performed target predictions using two steps; initially, the optimal complementarity between given mRNA and mature miRNA sequence was calculated using a weighted dynamic programming algorithm (Enright et al. 2003). This was then followed by estimation of free energy between mRNA and miRNA pair duplex structure using Vienna package (Lorenz et al. 2011). The network was then imported into Cytoscape software (Cytoscape_v3.8.2 URL: https://cytoscape.org/download.html) for further analysis (Shannon et al. 2003).

### 2.6 Computational validation of miRNA-Gene targets

The miRNA:mRNA targets obtained from prediction and network analysis were subjected to computational validation by submitting the target pairs to four different prediction algorithms, namely, RNA22v2 (Loher and Rigoutsos 2012), RNAhybrid (Krüger and Rehmsmeier 2006), miRmap (Vejnar et al. 2013), and TargetScan (Agarwal et al. 2015). The FASTA sequences of miRNAs and genes were downloaded from miRBase (URL: http://www.mirbase.org/; 02 March 2021; *Bos taurus*) and NCBI (URL: https://www.ncbi.nlm.nih.gov/gene/; 03 March 2021; *Bos taurus*), respectively. The genes and their miRNA target were subjected to all algorithms, and only those miRNA-Gene pairs that were validated by all the four algorithms were considered significant.

## 3. Result and Discussion

### 3.1 Lineage profile and functional annotation of DEGs among control and infected MECs

Differential gene expression carried out after processing with RMA algorithm and quantile normalization yielded 443 up-regulated and 92 down-regulated genes in *E.coli* infected MECs, whereas only 115 up-regulated and 16 down-regulated genes were observed in MECs infected with *S.aureus*. Correlation analysis using Z-score generated lineage profile for both *E.coli and S.aureus* infected MECs as represented in Fig 2 (a) and (b), respectively. The samples of *E.coli* and *S.aureus* had a higher Z-score for mammary gland, placenta, uterus and macrophages. In contrast, negative Z-scores were obtained for liver, spleen, lungs, stomach and neurons, thus indicating that the genes expressed in our samples indeed had mammary tissue expression lineage.

**Fig. 2:**
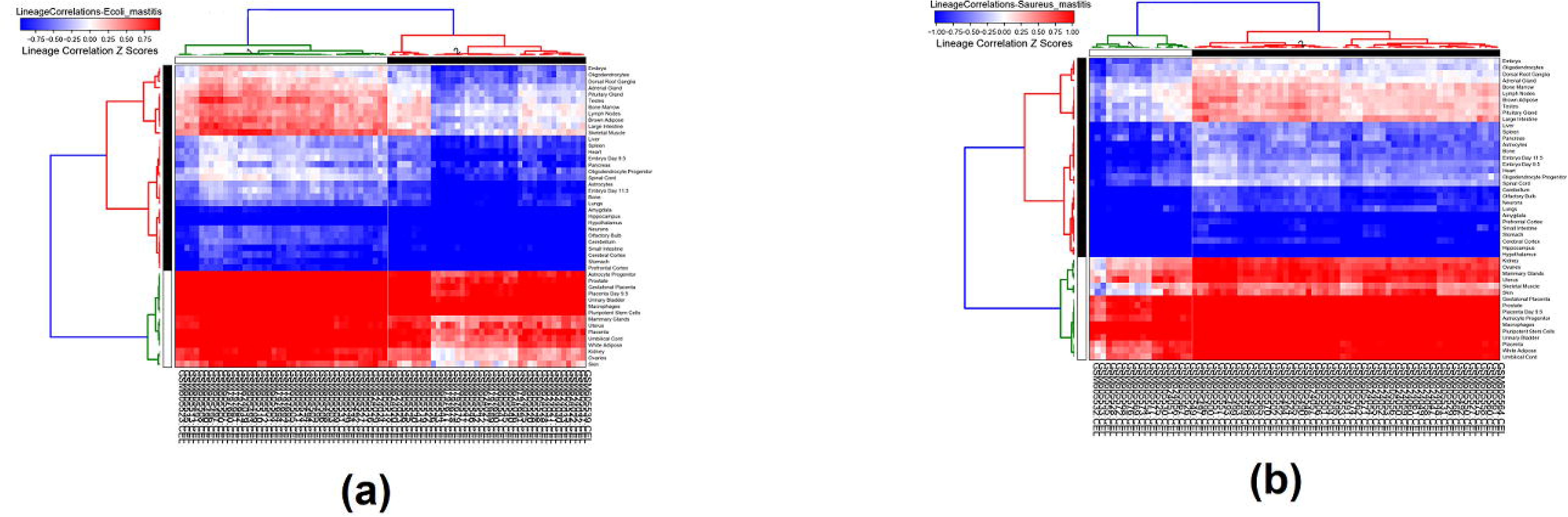
Z-score based lineage correlation profile of protein coding genes expressed in *E.coli* (a) and *S.aureus* (b) infected MECs.

Heatmap constructed from correlation analysis, using Z-score for functional gene ontology analysis, provided immune related pathway enriched by up-regulated genes of *E.coli* dataset (Fig 3 (a)). Type-1 helper 1 type immune response, cellular response to lipopolysaccharide, positive regulation of cytokine production and defense response were the ontology enriched in these samples. These results are consistent with a previous study on clinical mastitis, which reported increased cytokine production and lipopolysaccharide binding proteins such as IL-1B, IL-10 and heptoglobin in mastitis infected animals (Wenz et al. 2010). The major up-regulated genes in *E.coli* mastitis have been linked to induction and regulation of immune reactions and inflammatory responses (Buitenhuis et al. 2011). On the other hand, down-regulated genes in *E.coli* infected mastitis had their functional ontology for metabolic processes (Fig 3 (b)). Process like regulation of epithelial cell proliferation, developmental process and regulation of lipid metabolic process were among the enriched ontology. However, the z-score for these processes were not consistent in the analyzed cluster of down-regulated genes, which might suggest low credibility of their use in developing molecular markers of mastitis.

**Fig. 3:**
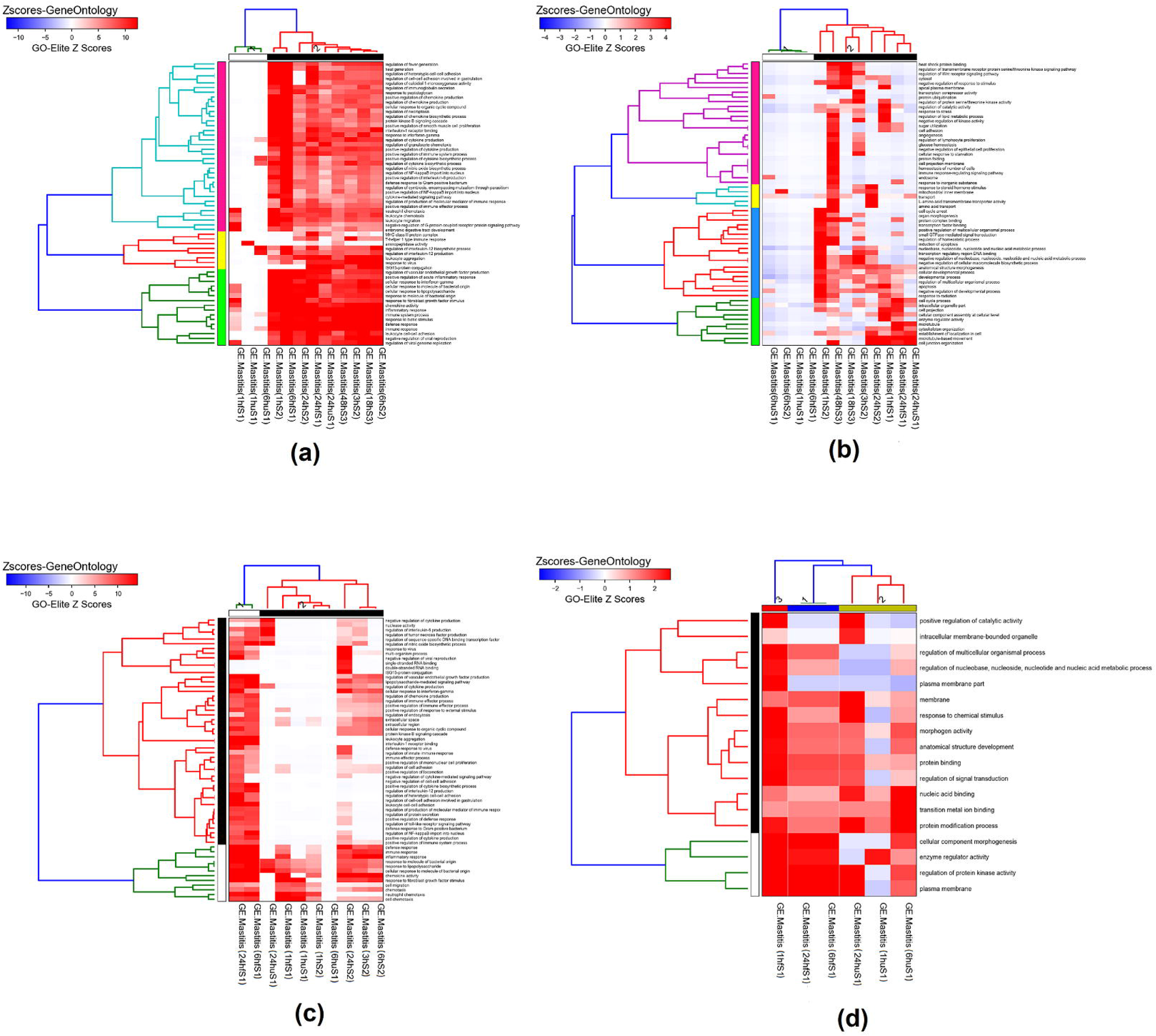
Functional ontology of (a) up-regulated and (b) down-regulated genes in *E.coli* and (c) up-regulated and (d) down-regulated genes in *S.aureus* infected MECs.

In *S.aureus* datasets, the gene ontology enriched by up-regulated genes (Fig 3(c)) included regulation of processes related to immune response, positive regulation of cytokine production, inflammatory responses and cellular response to molecules of bacterial origin; however, these ontology were not significantly enriched in samples taken 1 and 6 hour after infection especially in animals with unfavourable quantitative trait loci. On the other hand, the ontology enriched by down-regulated genes in *S.aureus* infected samples (Fig 3(d)) mainly belonged to cellular processes, as was observed with *E.coli* infected samples. Ontologies such as protein modification process, morphogen activity, transition metal ion binding and protein binding were the functionally enriched ontology for down-regulated genes in *S.aureus* gene expression data.

### 3.2 GSEA of DEGs among control and infected MECs

The GSEA of DEGs among control and *E.coli* infected MECs revealed that NOD-like receptor signalling pathway, Cytokine-cytokine interaction, NF-Kappa beta signalling pathway and Toll-like receptor signalling pathway were affected in *E.coli* infected MECs. All these pathways were related to immune reactions. NF-Kappa B signalling pathway (Fig 4A) had an enrichment score of 2.75 and the up-regulated protein coding genes include BAFF, ICAM that are found to responsible for B-Cell and T-Cell development. Also CD14, CD40, IL-1B and TNF- α were the other genes that were found to be significantly up-regulated in *E.coli* infected mastitis. NF-Kappa B signalling pathway has been previously been studied for its role in regulation of milk production wherein increased level of NF-Kappa B in epithelial cells was found to drastically reduce the milk production in mammary gland of bovines (Connelly et al. 2010). Another pathway enriched by the differential expressed genes of *E.coli* infected mastitis dataset that is linked to immune regulation was Toll-like receptor signalling pathway (Fig 4B). In Toll-like receptor signalling pathway, both TLR-2 and TLR-4 were found to be up-regulated, which are capable of inducing ubiquitin-mediated proteolysis, chemotatic and chemokine effects leading to inflammatory response that can be deduced as a host epithelial cell reaction to the presence infectious microbe. The cytokine up regulation in pathways were found to be instigated by immune response in bovine in response to various pattern recognition receptors in infected cells (Bhattarai et al. 2018).

**Fig. 4:**
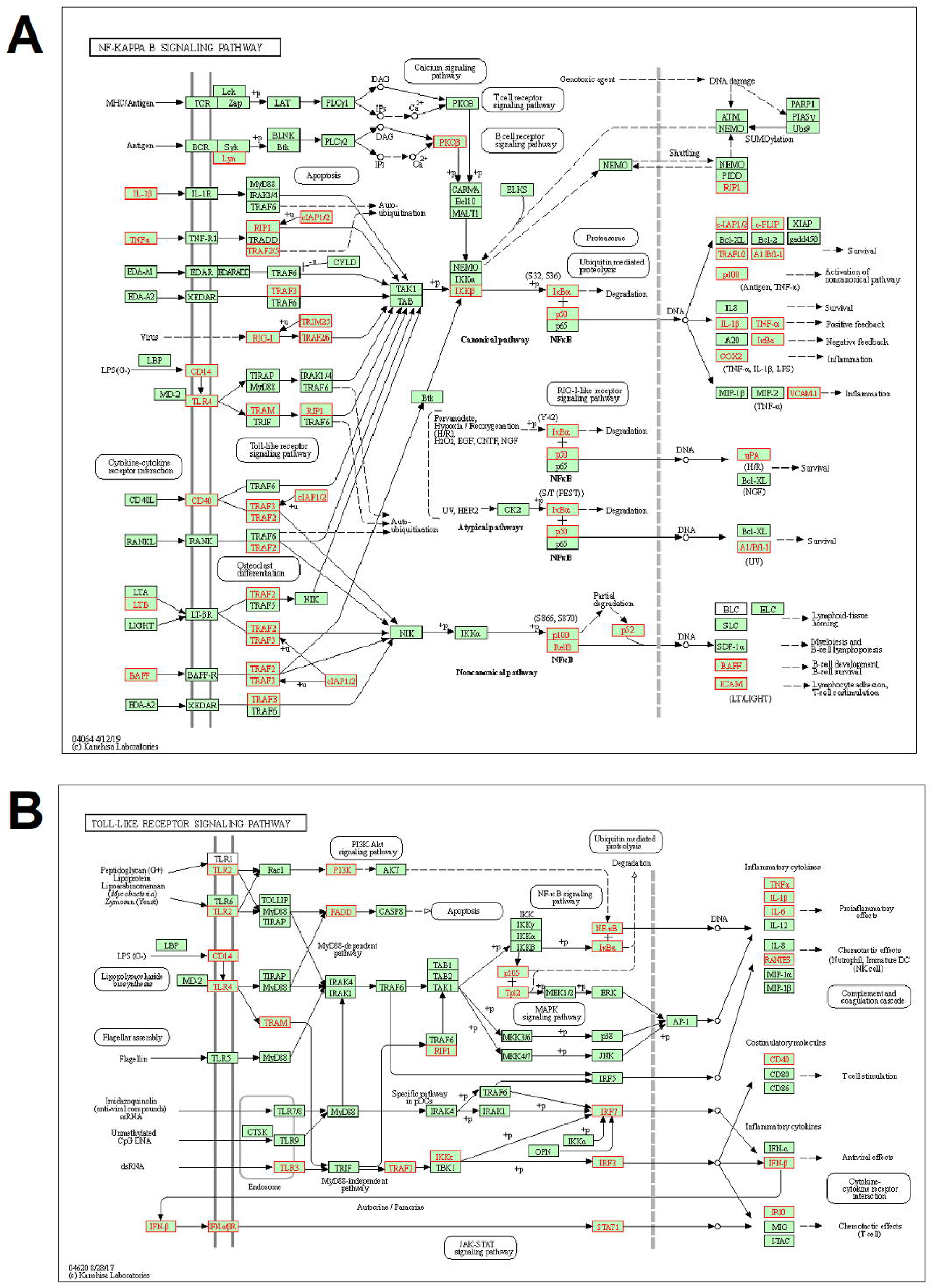
Pathways enriched by significant DEGs of E.coli infected MECs include (A) NF-Kappa B signalling pathway; (B) Toll-like receptor signalling pathway. (Genes highlighted in red are up-regulated during *E.coli* infection).

Similar to *E.coli* infected MECs, the pathways enriched by DEGs in *S.aureus* infected MECs also included TNF-signalling pathway, NF-kappa B signalling pathway, Toll-like receptor signalling pathway and other immune regulation pathways. NF- Kappa B signalling pathway was positively enriched with a score of 2.9 (Fig 5A) and was found to be involved in immunogenic reaction of host against *S.aureus* infection. Genes such as TNF-α, IL-1β, CD14, CD40, ICAM, and VCAM1 were found to be up-regulated in case of *S.aureus* infection. An interesting observation in the up-regulated genes is that, unlike *E.coli* infection wherein both humoral and cellular response appears to be induced, only T-Cells stimulation appears to be activated in *S.aureus* infection indicating there may be involvement of only cellular immunity in *S.aureus* infection during initial infection. Toll-like receptor signalling pathway (Fig 5B) was another pathway that was significantly enriched in *S.aureus* infection. Interestingly, in *S.aureus* infection, only TLR2 was activated whereas in case of *E.coli* infection both TLR2 and TLR4 were found to be activated during infection. This correlated with a previous study aimed at quantifying TLR expression during mastitis wherein the subclinical nature of mastitis was attributed to down regulation of soluble TLR2 production by serum constituents hindering the immune reactions (Fu et al. 2013). The up-regulated genes in this pathway also included pro-inflammatory genes, genes responsible for chemotatic effects and genes involved in T-Cell stimulation.

**Fig. 5:**
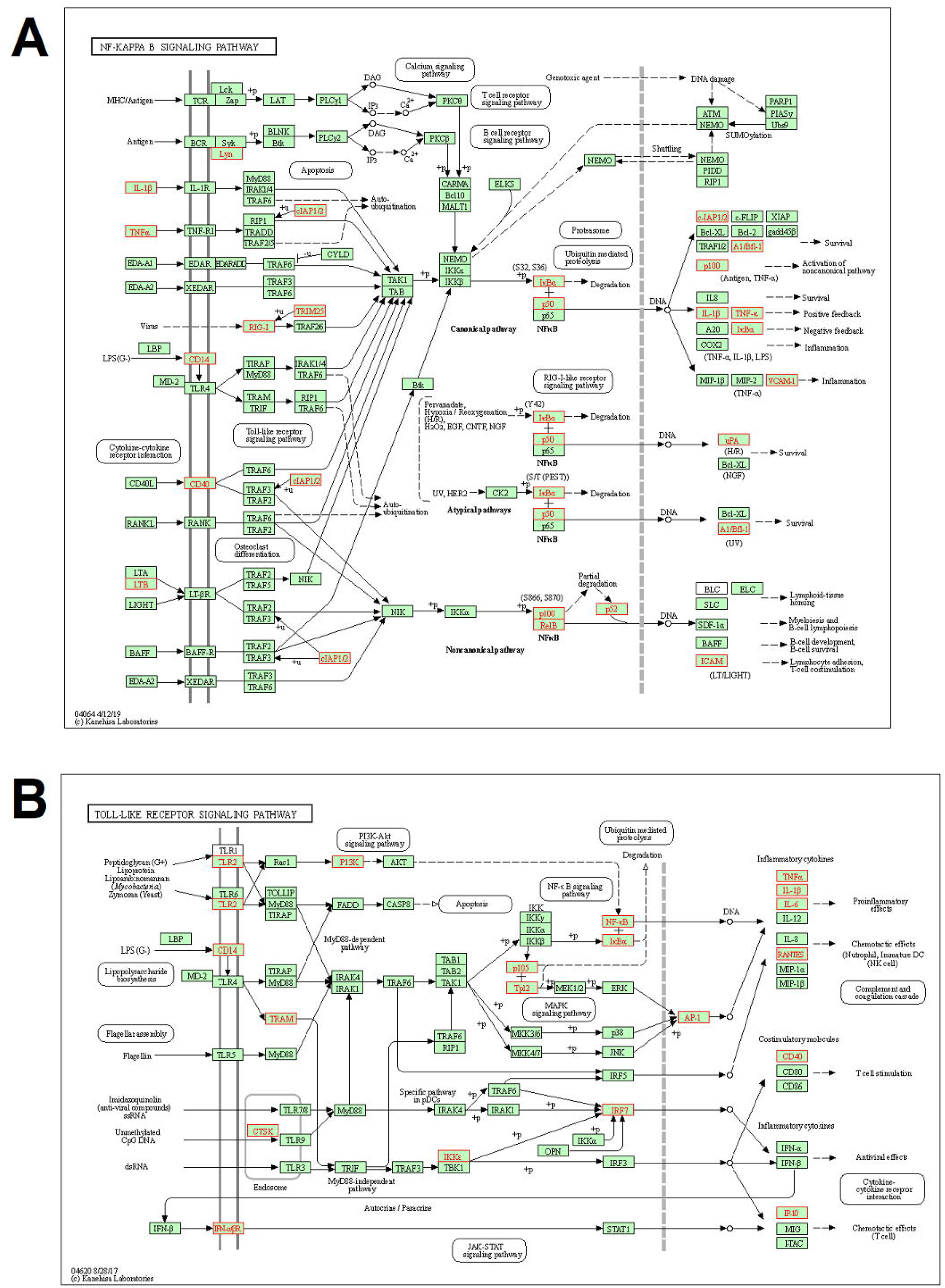
Pathways enriched by significant DEGs of *S.aureus* infected MECs include (A) NF-Kappa B signalling pathway; (B) Toll-like receptor signalling pathway. (Genes highlighted in red are up-regulated during *S.aureus* infection).

### 3.3 Meta-analysis of DEGs of *E.coli* and *S.aureus* samples

#### 3.3.1 Meta-analysis by combined p-value method

Meta-analysis of DEGs among control and infected MECs was conducted using two statistical methods. The first method was using combined p-value method. The top 10 up-regulated and down- regulated genes of *E.coli* infected MECs are presented in Table 1; Volcano plot obtained after meta-analysis is represented in Fig 6A. Several up-regulated genes were found to be part of chemokine signalling pathway and included mastitis related genes like CXCL2, CCL20, IL8, and NFKBIZ. However, in *S.aureus* infected MECs, the meta-analysis of DEGs resulted in considerably less number of significant up-regulated and down-regulated genes. The volcano plot obtained after meta-analysis for *S.aureus* infected MECs (Fig 6B) showed that p-values after analysis were not significant, as was seen in *E.coli* datasets. Top up-regulated and down-regulated genes for *S.aureus* datasets are given in Table 1, which included CXCL2, CYP1A1 and CXCL5 genes.

**Table 1:**
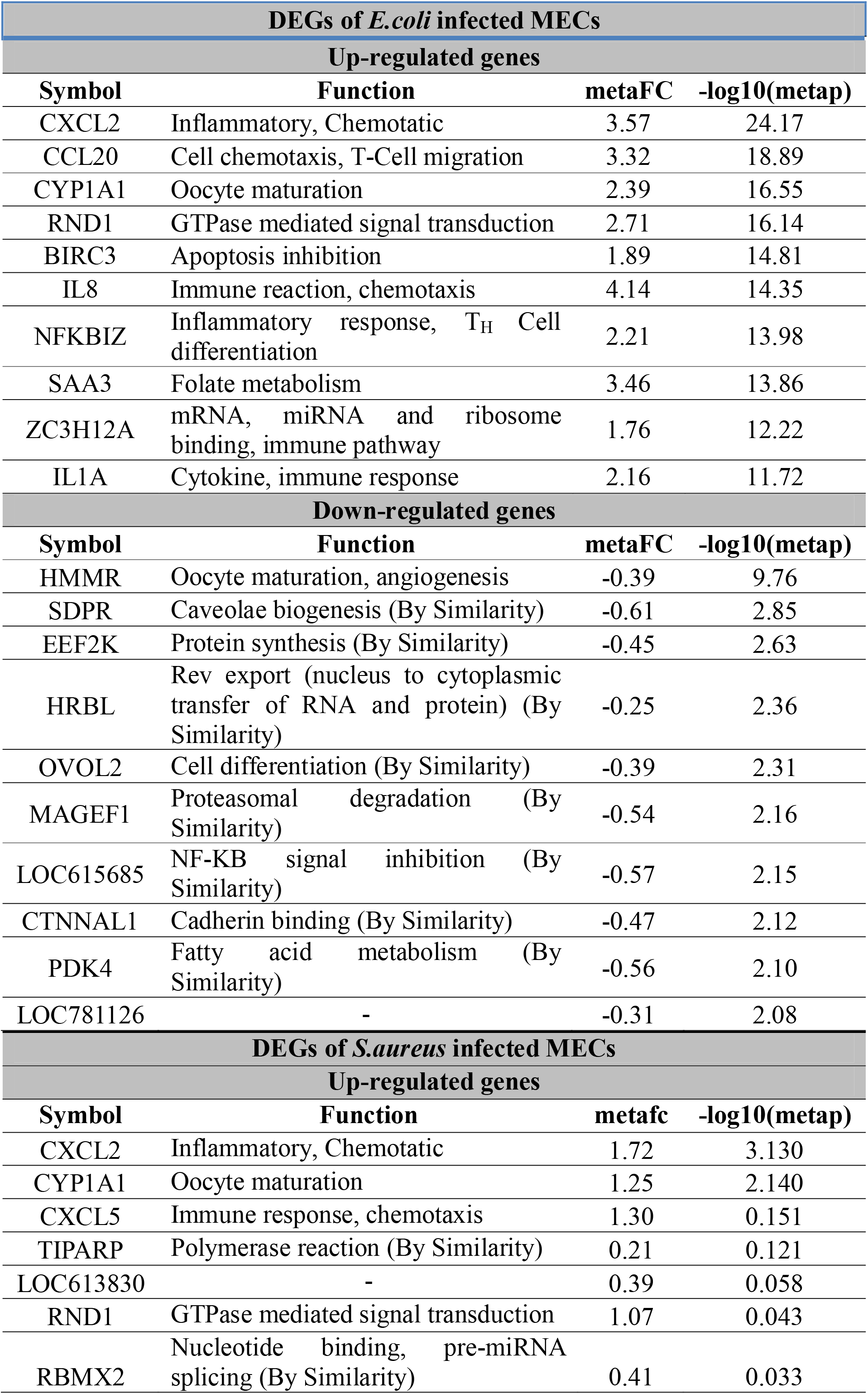
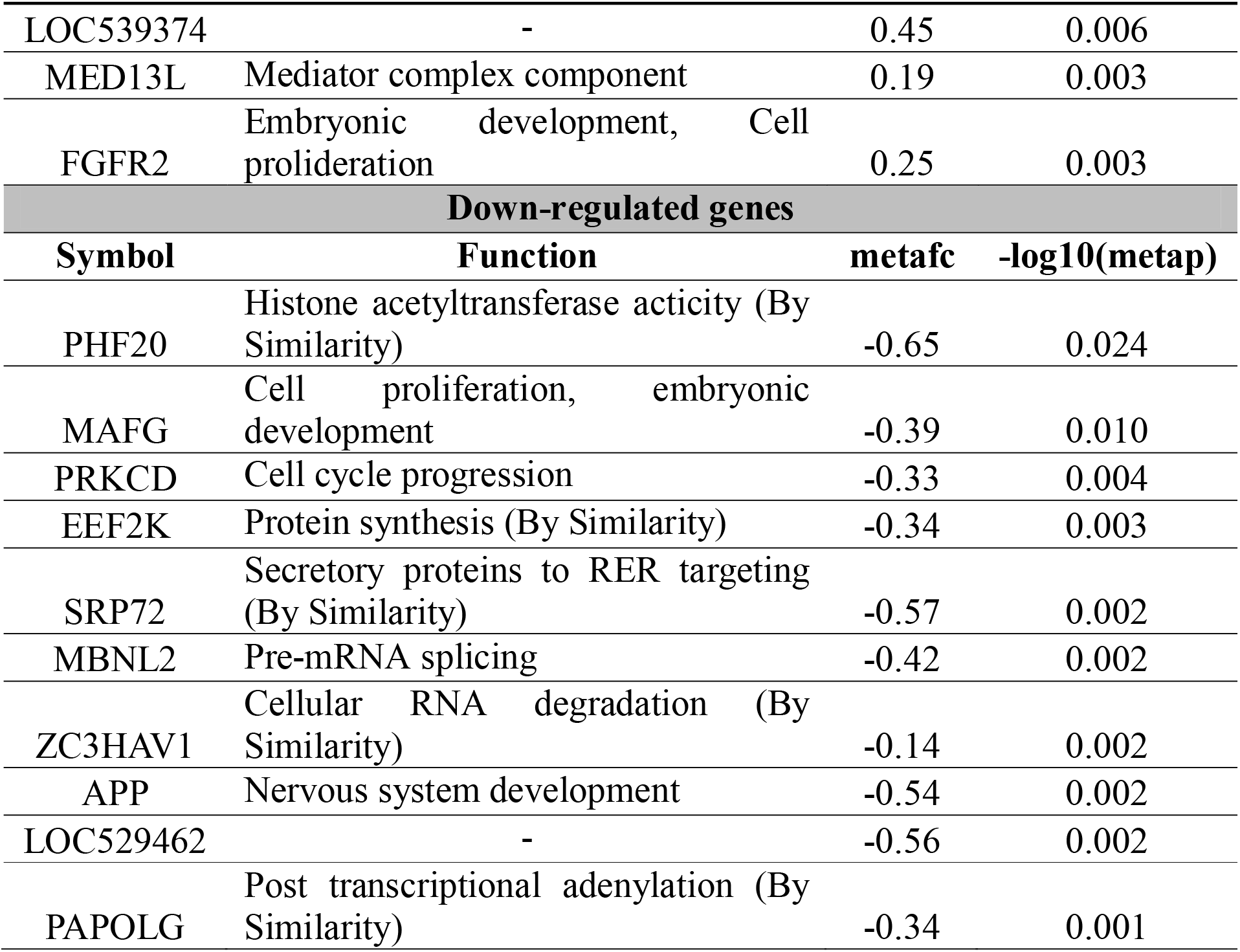
Significant differentially expressed genes in *E.coli* and *S.aureus* datasets using combined p-value method

**Fig 6:**
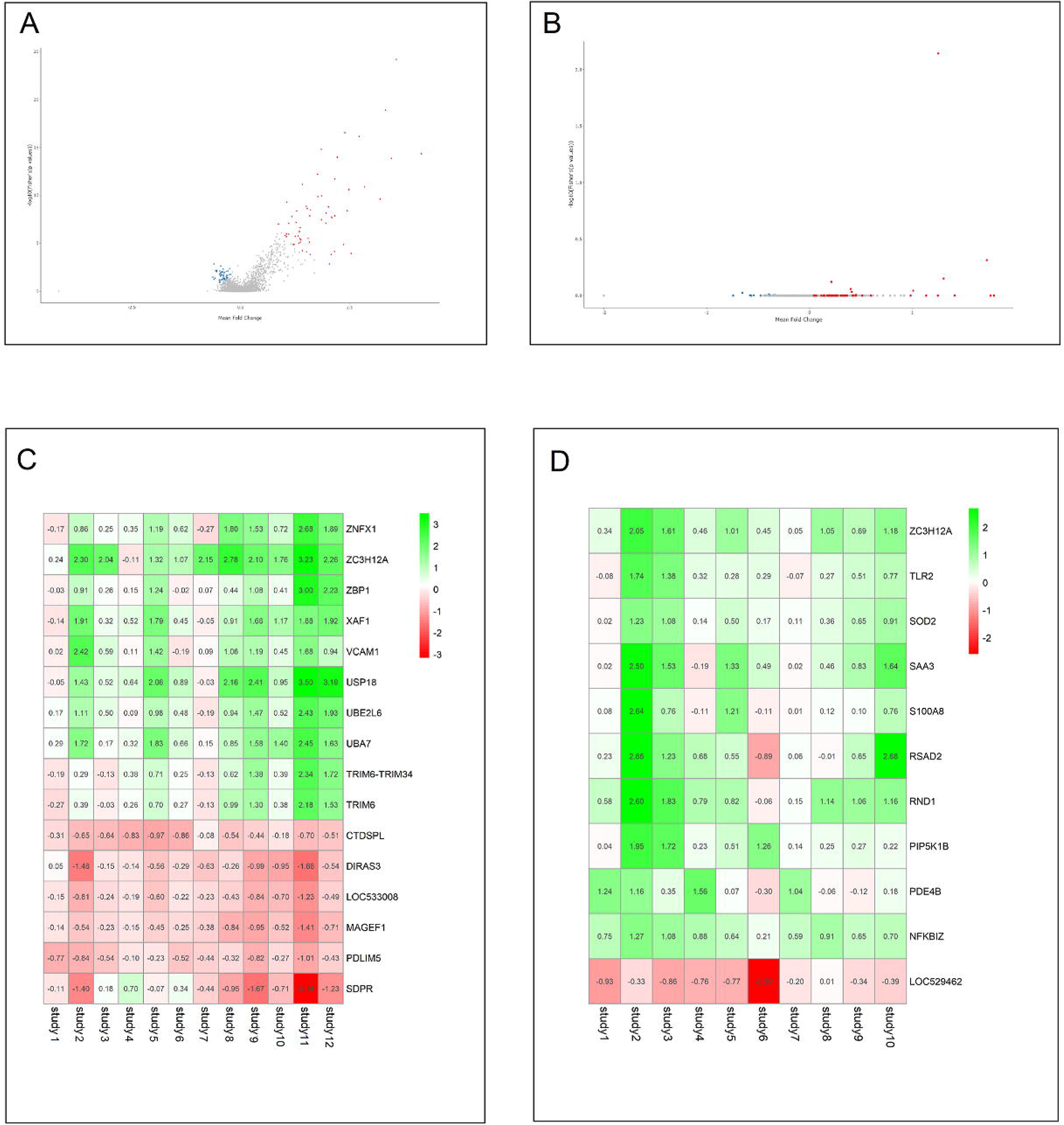
Volcano plot for differentially expressed genes from meta-analysis of *E.coli* (A) and *S.aureus* (B) infected MECs; Significant up- and down-regulated genes obtained from robust rank aggregation of datasets from *E.coli* (C) and *S.aureus* (D) infected MECs.

#### 3.3.2 Meta-analysis by RRA

The second method used for meta-analysis of DEGs was RRA method. The significant DEGs for *E.coli* datasets is as shown in Fig. 6C and their corresponding p-values are presented in Table 2, respectively. The significant DEGs of *S.aureus* datasets after meta-analysis using RRA method showed 31 up-regulated and 1 down-regulated genes. The top DEGs with their expression values are presented in Fig 6D, and the corresponding log fold changes with their p-value after evaluation are presented in Table 2, respectively. Interestingly, ZC3H12A gene was found to be up-regulated in both *E.coli* and *S.aureus* induced mastitis infection and ZC3H12A has previously been studied for its control in stability of inflammatory genes (Matsushita et al. 2009).

**Table 2:**
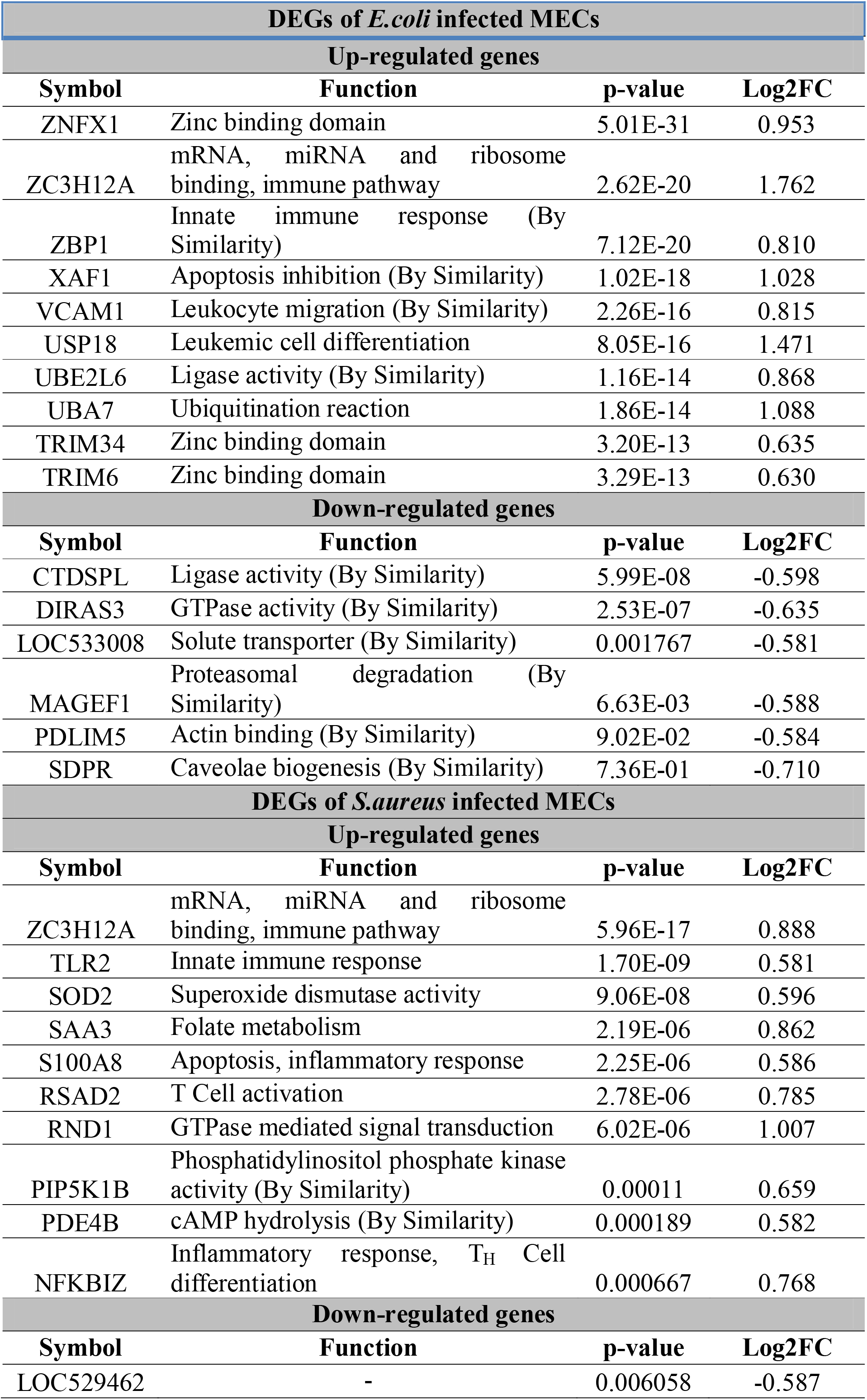
Significant differentially expressed genes in *E.coli* and S.aureus datasets using robust rank aggregation

#### 3.3.3 Identification of significant DEGs after meta-analysis

Genes significant from the p-value and RRA methods were screened by extracting the intersection of obtained gene sets for up-regulated and down-regulated genes of *E.coli* and *S.aureus*, respectively. After analysis and comparison, 43 genes were found to be common up-regulated genes. Five genes were found to be the significantly down-regulated genes in *E.coli* (Fig 7(a) and 7(b)) whereas six common genes were found to be up-regulated and one common gene was found to be down-regulated in *S.aureus* infected MECs (Fig 7(c) and (d)). Table 3 provide the common significant genes obtained of *E.coli* and *S.aureus* infected mastitis samples, respectively.

**Fig. 7:**
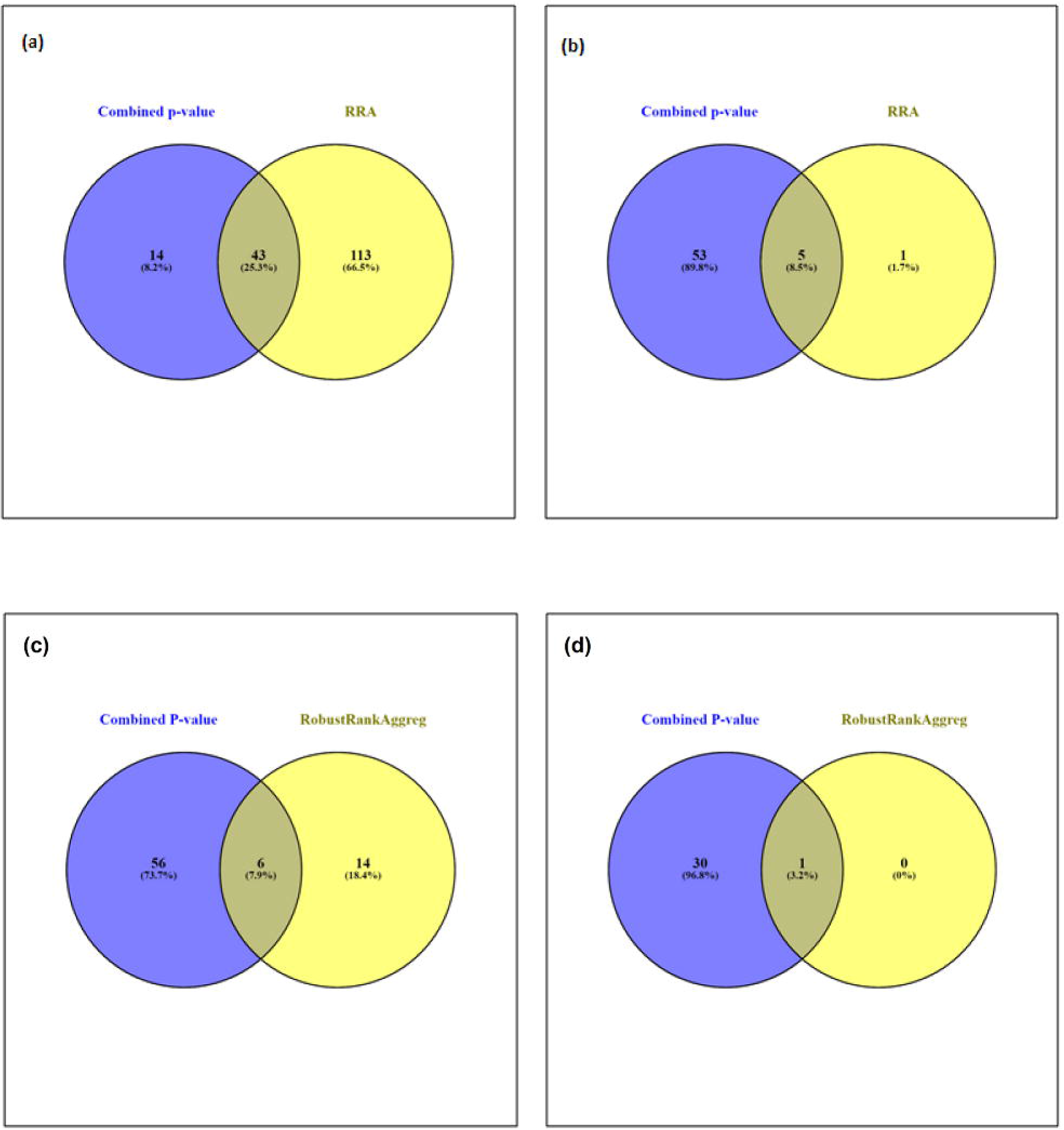
Venn diagram representing intersection of significant DEGs (a) Upregulated and (b) downregulated in *E.coli* and (c) Upregulated and (d) Downregulated in *S.aureus* infected MECs.

**Table 3:** Common significant DEGs in *E.coli* and *S,aureus* datasets

| DEGs of *E.coli* infected MECs |  |
| --- | --- |
| Up-regulated genes |  |
| Symbol | Function |
| IL8 | Immune reaction, chemotaxis |
| SAA3 | Folate metabolism |
| RND1 | GTPase mediated signal transduction |
| NFKBIZ | Inflammatory response, T<sub>H</sub> Cell differentiation |
| ZC3H12A | mRNA, miRNA and ribosome binding, immune pathway |
| IL1A | Cytokine, immune response |
| MRM1 | Methyltransferase reaction |
| SOD2 | Superoxide dismutase activity |
| IL1B | Cytokine activity |
| NFKBIA | NF-KB/REL dimer formation inhibition |
| IL6 | Cytokine activity |
| NOS2 | Nitric oxide synthase |
| HP | Antioxidant activity |
| RSAD2 | T Cell activation |
| MX2 | GTP binding (immune process) |
| LOC507402 | - |
| MX1 | GTP binding (immune process) |
| ISG15 | Binding to integrin (immune response) |
| S100A9 | Antioxidant activity (inflammatory response) |
| MAP3K8 | TLR4 signalling pathway (By Similarity) |
| TNFSF13B | B Cell activation |
| TLR2 | Innate immune response |
| LTF | Lipopolysaccharide binding, kinase activator |
| TNIP1 | Inhibits NF-KB activation (By Similarity) |
| LOC518495 | - |
| IRF1 | Transcription regulation (immune related) |
| ISG20 | Exoribonuclease activity |
| IFNAR2 | Interferon receptor (By Similarity) |
| S100A8 | Apoptosis, inflammatory response |
| RTP4 | Cell surface expression of GPCR (By Similarity) |
| PIP5K1B | Phosphatidylinositol phosphate kinase activity (By Similarity) |
| USP18 | Leukemic cell differentiation |
| LGP2 | Antiviral signalling |
| LOC517354 | Chemotatic activity |
| IFIH1 | Innate immune receptor (By Similarity) |
| IFIT2 | Antiviral activity |
| IL18BP | Pro-inflammatory inhibition |
| MGC143170 | NF-KB signalling |
| ISG12(B) | - |
| LOC512486 | Innate immunity |
| KLK10 | Serine endopeptidase activity |
| LTB | Immune response |
| RNF19B | T Cell, NK Cell cytolytic activity |
| **Down-regulated genes** |  |
| **Symbol** | **Function** |
| DIRAS3 | GTPase activity (By Similarity) |
| CTDSPL | Ligase activity (By Similarity) |
| SDPR | Caveolae biogenesis (By Similarity) |
| MAGEF1 | Proteasomal degradation (By Similarity) |
| LOC533008 | Solute transporter (By Similarity) |
| **DEGs of *S.aureus* infected MECs** |  |
| **Up-regulated genes** |  |
| **Symbol** | **Function** |
| SAA3 | Folate metabolism |
| IL8 | Immune reaction, chemotaxis |
| NFKBIZ | Inflammatory response, T<sub>H</sub> Cell differentiation |
| RND1 | GTPase mediated signal transduction |
| MAP3K8 | TLR4 signalling pathway (By Similarity) |
| ZC3H12A | mRNA, miRNA and ribosome binding, immune pathway |
| **Down-regulated genes** |  |
| **Symbol** | **Function** |
| LOC529462 | - |

### 3.4 miRNA-Gene network of significant DEGs

The miRNA-Gene network constructed for *E.coli* DEGs had 158 nodes and 190 edges and consisted of both up-regulated and down-regulated genes (Fig 8A). The miRNA-Gene pairs present in the repository are either predicted targets from miRanda or experimentally validated target sets. Gene-miRNA network target obtained for *S.aureus* DEGs the final network had 37 nodes and 34 edges as shown in Fig 8B. Common gene markers, having miRNA targets from both datasets, included the ZC3H12A, RND1 and MAP3K8, which were up-regulated and may be potential markers of mastitis. Gene with miRNA targets exclusive to *E.coli* infection included RSAD2, IRF1, KLK10, MRM1, IFNAR2, TLR2, CTDSPL, RNF19B, LTB and SDPR. These genes may probably be used for distinguishing *E.coli* infection during mastitis.

**Fig 8:**
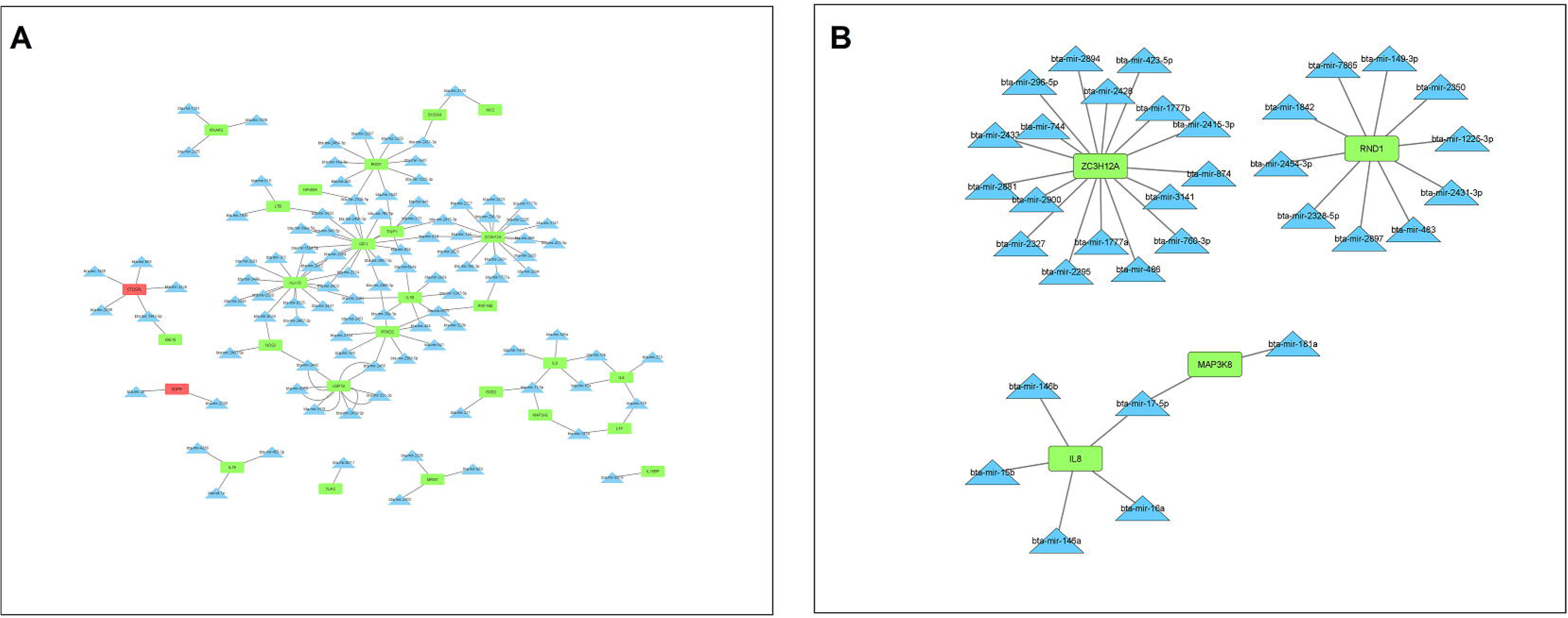
The miRNA-Gene target network in *E.coli* (A) and *S.aureus* (B) infected MECs.

### 3.5 Computational validation of miRNA-Gene targets

All the significant targets for *E.coli and S.aureus* infected MECs are presented in Table 4 and Table 5, respectively. Among them, target pairs bta-miR-1777b:ZC3H12A, bta-miR-2900: ZC3H12A, bta-miR-2881: ZC3H12A, bta-miR-3141: ZC3H12A, bta-miR-149-3p: RND1; bta-miR-2328-5p: RND1; bta-miR-181a: MAP3K8 were particularly more enticing as these miRNA- gene pairs were identified in both *E.coli* and *S.aureus* mastitis. Studies have shown that genes targeted by miRNA in mastitis were primarily involved in immune or inflammatory pathway (Lai et al. 2017). MicroRNA targeting genes involved in intracellular protein transport and protein binding like bta-miR-185 and bta-miR-378 were previously reported to be up-regulated in *S.aureus* infected cow milk (Ma et al. 2019). Studies have also reported lowered expression of bta-miR-181a to regulate the TLRs expression in *S.uberis* infected MECs (Naeem et al. 2012), which correlates with the results of the present study that bta-miR-181a targeted the MAP3K8 expression. Further characterization and abundance of these miRNAs and their targeted gene counterparts in MECs may thus, qualify them as potential molecular markers of mastitis.

**Table 4:** Computationally validated miRNA-Gene targets of significant *E.coli* DEGs

| miRNA | Gene | RNA22<br>Kcal/mol | RNAhybrid<br>(minimum free<br>energy) | miRmap<br>score | Site<br>type | Context<br>++ score<br>% | Context<br>++ score | Context ++<br>score<br>Weighted | Branch<br>length<br>conserved |
| --- | --- | --- | --- | --- | --- | --- | --- | --- | --- |
| bta-miR-6525 | RSAD2 | -22 | -28.4 | 94.1582 | 8mer | 98 | -0.39 | -0.39 | 0.439 |
| bta-miR-2465 | RSAD2 | -26.2 | -32.5 | 96.0599 | 7mer-m8 | 91 | -0.21 | -0.21 | 0 |
| bta-miR-2453 | RSAD2 | -20 | -30.2 | 93.5593 | 7mer-m8 | 92 | -0.2 | -0.2 | 0 |
| bta-miR-149-3p | RND1 | -25.5 | -35.8 | 91.9972 | 7mer-m8 | 44 | -0.02 | -0.02 | 0.338 |
| bta-miR-2328-5p | RND1 | -29.3 | -37.6 | 96.3532 | 7mer-m8 | 83 | -0.17 | -0.17 | 0 |
| bta-miR-2412 | IRF1 | -27.4 | -37 | 99.4617 | 7mer-m8 | 91 | -0.25 | -0.25 | 0 |
| bta-miR-2447 | KLK10 | -17.3 | -32.8 | 97.9702 | 7mer-m8 | 91 | -0.28 | -0.28 | 0 |
| bta-miR-2412 | KLK10 | -33 | -41.3 | 94.6849 | 7mer-m8 | 91 | -0.28 | -0.28 | 0 |
| bta-miR-2374 | KLK10 | -33.6 | -38.3 | 94.8621 | 7mer-m8 | 91 | -0.28 | -0.28 | 0 |
| bta-miR-1584-5p | KLK10 | -25.4 | -34.5 | 93.8459 | 7mer-m8 | 91 | -0.28 | -0.28 | 0 |
| bta-miR-2392 | KLK10 | -33.8 | -39.2 | 98.3607 | 7mer-m8 | 77 | -0.16 | -0.16 | 0 |
| bta-miR-2893 | KLK10 | -31.4 | -38.2 | 98.7386 | 7mer-m8 | 78 | -0.09 | -0.09 | 0 |
| bta-miR-2467-3p | KLK10 | -20.7 | -38.9 | 97.1999 | 7mer-m8 | 88 | -0.15 | -0.15 | 0 |
| bta-miR-2360 | KLK10 | -26 | -38.9 | 99.8289 | 7mer-m8 | 93 | -0.3 | -0.3 | 0 |
| bta-miR-2889 | IL1B | -30.2 | -35.8 | 93.5893 | 7mer-m8 | 78 | -0.19 | -0.19 | 0 |
| bta-miR-484 | IL1B | -15.7 | -33.1 | 95.9381 | 7mer-A1 | 83 | -0.12 | -0.12 | 0 |
| bta-miR-320b | IL1B | -21.3 | -30.4 | 95.0999 | 7mer-m8 | 96 | -0.27 | -0.27 | 0 |
| bta-miR-1777b | ZC3H12<br>A | -35.3 | -45.4 | 99.6471 | 7mer-m8 | 89 | -0.35 | -0.35 | 0 |
| bta-miR-2900 | ZC3H12<br>A | -28.9 | -36.9 | 99.5642 | 7mer-m8 | 90 | -0.39 | -0.39 | 0 |
| bta-miR-2881 | ZC3H12<br>A | -31.6 | -39.2 | 99.8310 | 7mer-m8 | 93 | -0.37 | -0.37 | 0 |
| bta-miR-3141 | ZC3H12<br>A | -22.2 | -35.6 | 99.2012 | 7mer-m8 | 92 | -0.32 | -0.32 | 0 |
| bta-miR-181a | MAP3K8 | -13.6 | -24.8 | 96.8592 | 7mer-m8 | 69 | -0.08 | -0.08 | 3.033 |
| bta-miR-2306 | MRM1 | -19.9 | -32.9 | 96.9702 | 7mer-m8 | 88 | -0.19 | -0.19 | 0 |
| bta-miR-2305 | IFNAR2 | -20.8 | -38 | 99.6536 | 8mer | 73 | -0.27 | -0.12 | 0 |
| bta-miR-1291 | IFNAR2 | -23.3 | -34.6 | 95.1727 | 7mer-m8 | 74 | -0.16 | -0.07 | 0 |
| bta-miR-6517 | TLR2 | -14.9 | -32.5 | 93.2993 | 7mer-A1 | 82 | -0.12 | -0.12 | 0 |
| bta-miR-2899 | CTDSPL | -16.2 | -34.7 | 97.1194 | 8mer | 91 | -0.32 | -0.25 | 0 |
| bta-miR-1343-5p | CTDSPL | -14.4 | -36.2 | 96.6853 | 7mer-m8 | 75 | -0.09 | -0.09 | 0 |
| bta-miR-658 | CTDSPL | -22.4 | -35 | 94.6540 | 7mer-m8 | 82 | -0.24 | -0.24 | 0 |
| bta-miR-1777a | RNF19B | -31.4 | -44.1 | 98.6908 | 7mer-m8 | 89 | -0.27 | -0.01 | 0 |
| bta-miR-6525 | RNF19B | -16.2 | -28.9 | 95.5793 | 7mer-m8 | 50 | -0.05 | 0 | 0 |
| bta-miR-615 | LTF | -32.9 | -43.7 | 96.8394 | 8mer | 97 | -0.4 | -0.4 | 0.573 |
| bta-miR-2290 | SDPR | -19.1 | -32.4 | 95.4715 | 7mer-m8 | 94 | -0.18 | -0.18 | 0 |

**Table 5:** Computationally validated miRNA-Gene targets of significant *S.aureus* DEGs

| miRNA | Gene | RNA22<br>Kcal/mol | RNAhybrid<br>(minimum free<br>energy) | miRmap<br>score | Site<br>type | Context<br>++<br>score % | Context<br>++<br>score | Context ++<br>score<br>weighted | Branch<br>length<br>conserved |
| --- | --- | --- | --- | --- | --- | --- | --- | --- | --- |
| bta-miR-1777b | ZC3H12<br>A | -35.3 | -45.4 | 99.6471 | 7mer-m8 | 89 | -0.35 | -0.35 | 0 |
| bta-miR-2900 | ZC3H12<br>A | -28.9 | -36.9 | 99.5642 | 7mer-m8 | 90 | -0.39 | -0.39 | 0 |
| bta-miR-2881 | ZC3H12<br>A | -31.6 | -39.2 | 99.8310 | 7mer-m8 | 93 | -0.37 | -0.37 | 0 |
| bta-miR-3141 | ZC3H12<br>A | -22.2 | -35.6 | 99.2012 | 7mer-m8 | 92 | -0.32 | -0.32 | 0 |
| bta-miR-149-3p | RND1 | -25.5 | -35.8 | 91.9972 | 7mer-m8 | 44 | -0.02 | -0.02 | 0.338 |
| bta-miR-2328-5p | RND1 | -29.3 | -37.6 | 96.3532 | 7mer-m8 | 83 | -0.17 | -0.17 | 0 |
| bta-miR-181a | MAP3K8 | -13.6 | -24.8 | 96.8592 | 7mer-m8 | 69 | -0.08 | -0.08 | 3.033 |

## 5. Conclusion

In the present study, meta-analysis of high throughput transcriptomic data of MECs was carried out to predict the molecular markers of *E.coli* and *S.aureus* infected mastitis by combined p-value and robust rank aggregation. The combined use of two methods provided the top perturbed genes in mastitis infection, which could be potential molecular markers of mastitis. The combined p-value and RRA avoided biases produced by different experimental setup whereas RRA reduced the type I errors. Immune related NF-Kappa B signalling pathway and Toll like receptor signalling pathway were primarily affected in mastitis. However, *S.aureus* infection lead to activation of TLR2 only whereas *E.coli* activated both TLR2 and TLR4 during the course of infection. The ZC3H12A, RND1 and MAP3K8 were found to be perturbed in both *E.coli* and *S.aureus* infection whereas significantly increased expression of IFNAR2, TLR2, CTDSPL, RNF19B, LTB, SDPR, RSAD2, IRF1, KLK10 and MRM1 genes were observed in *E.coli* infection alone and may serve molecular marker to distinguish from *S.aureus* infection. In future, NF-Kappa B signalling pathway and Toll like receptor signalling pathway may also be explored as potential targets for drug development against prevention and treatment of mastitis.

